# Learning prediction error neurons in a canonical interneuron circuit

**DOI:** 10.1101/2020.02.27.968776

**Authors:** Loreen Hertäg, Henning Sprekeler

**Affiliations:** Technical University Berlin & BCCN Berlin

## Abstract

Sensory systems constantly compare external sensory information with internally generated predictions. While neural hallmarks of prediction errors have been found throughout the brain, the circuit-level mechanisms that underlie their computation are still largely unknown. Here, we show that a well-orchestrated interplay of three interneuron types shapes the development and refinement of negative prediction-error neurons in a computational model of mouse primary visual cortex. By balancing excitation and inhibition in multiple pathways, experience-dependent inhibitory plasticity can generate different variants of prediction-error circuits, which can be distinguished by simulated optogenetic experiments. The experience-dependence of the model circuit is consistent with that of negative prediction-error circuits in layer 2/3 of mouse primary visual cortex. Our model makes a range of testable predictions that may shed light on the circuitry underlying the neural computation of prediction errors.

## Introduction

Changes in sensory inputs can arise from changes in our environment, but also from our own movements. When you walk through a room full of people, your perspective changes over time, and you will experience a global visual flow. Superimposed on this global change are local changes generated by the movements of the people around you. An essential task of sensory perception is to disentangle these different origins of sensory inputs, because the appropriate behavioral responses to environmental and to self-generated changes are often different. Am I approaching a person or is she approaching me?

A common assumption is that perceptual systems subtract from the sensory data an internal prediction ^1–6^, which is calculated from an efference copy of the motor signals our brain has issued. Changes in the external world then take the form of mismatches – or prediction errors – between internal predictions and sensory data^7^. This comparison requires an accurate prediction system that adapts to ongoing changes in the environment or in behavior. An efficient way to ensure a flexible adaptation is to render the prediction circuits experience-dependent by minimizing prediction errors ^8^.

Neural hallmarks of prediction errors are found throughout the brain. Dopaminergic neurons in the basal ganglia and the striatum^9^ encode a reward prediction error (mismatch between expected and received reward), and subsets of neurons in visual cortex ^10,11^, auditory cortex ^12,13^ and barrel cortex ^14^ code for a mismatch between feedback and feedforward information.

While neural correlates of prediction errors have been found broadly, the circuit level mechanisms that underlie their computation are poorly understood. Given that prediction errors involve a subtraction of expectations from sensory data, the relevant circuits likely involve both excitatory and inhibitory pathways ^11^. Negative prediction-error (nPE) neurons, which are activated only when sensory signals are weaker than predicted, are likely to receive excitatory predictions counterbalanced by inhibitory sensory signals. Conversely, positive prediction-error (pPE) neurons, which respond only when sensory signals exceed the internal prediction, could receive excitatory sensory signals counterbalanced by inhibitory predictions. How the complex inhibitory circuits of the cortex ^15–18^ support the computations of these prediction errors is not resolved and neither are the activity-dependent forms of plasticity that would allow these circuits to refine the prediction machine.

For prediction-error neurons, fully predicted sensory signals should cancel with the internal prediction and hence trigger no response. We therefore hypothesized that an experience-dependent formation and refinement of prediction-error circuits can be achieved by balancing excitation and inhibition in an activity-dependent way. Using a computational model comprised of excitatory pyramidal cells and three types of inhibitory interneurons, we show that nPE neurons can be learned by inhibitory synaptic plasticity rules that balance excitation and inhibition in principal cells. We find that the circuit shows a similar experience dependence as observed in V1 ^11^. Depending on which interneuron classes receive motor predictions and which receive sensory signals, the plasticity rules shape different, fully functional variants of the prediction circuit. Using simulated optogenetic experiments, we show that these variants have identifiable fingerprints in their reaction to optogenetic activation or inactivation of different interneuron classes. Finally, we demonstrate that the inhibitory prediction circuits can be learned by biologically plausible forms of homeostatic inhibitory synaptic plasticity, which only rely on local information available at the synapses.

## Results

We studied a rate-based network model of layer 2/3 of rodent V1 to investigate how negative prediction-error (nPE) neurons develop. The model includes excitatory pyramidal cells (PCs) as well as inhibitory parvalbumin-expressing (PV), somatostatin-expressing (SOM) and vasoactive intestinal peptide-expressing (VIP) interneurons (Fig. 1a). All neurons in the model receive excitatory background input that ensures reasonable baseline activities in the absence of visual input and motor-related internal predictions (“baseline”). A subset of inhibitory synapses – chosen based on a mathematical analysis – are subject to experience-dependent plasticity, which homeostatically controls the firing rate of PCs by balancing excitation and inhibition ^19^(see Methods and Fig. 1a). We stimulated the network with time-varying external inputs that represent visual stimuli and motor-related internal predictions (Fig. 1a,b). We reasoned that during natural conditions, movements lead to sensory inputs that are fully predicted by internal motor commands (“feedback phase”^11^), while unexpected external changes in the environment should generate unpredicted sensory signals (“playback phase”^11^). Situations in which internal motor commands are not accompanied by corresponding sensory signals should be rare (“feedback mismatch phase”^11^). During plasticity, we therefore stimulated the circuit with a sequence consisting of feedback and playback phases (“quasi-natural training”, Fig. 1b).

**Figure 1.**
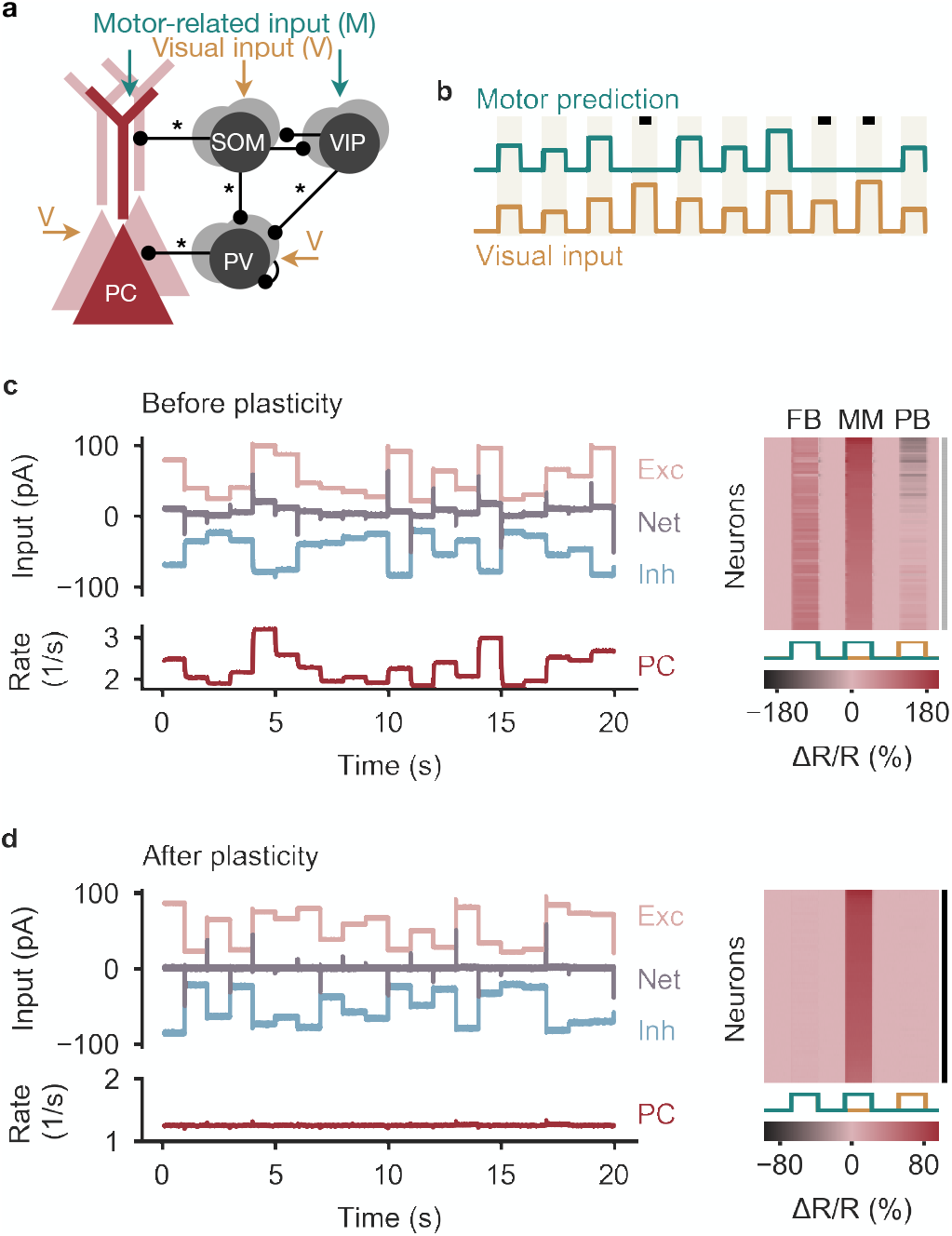
Balancing excitation and inhibition gives rise to negative prediction-error neurons. **(a)** Network model with excitatory PCs and inhibitory PV, SOM and VIP neurons. Connections from PCs onto inhibitory neurons not shown for the sake of clarity. Somatic compartment of PCs, SOM and PV neurons receive visual input, apical dendrites of PCs and VIP neurons receive a motor-related prediction thereof. Connections marked with an asterisk undergo experience-dependent plasticity. **(b)** During plasticity, the network is exposed to a sequence of feedback (coupled sensorimotor experience) and playback phases (black square, visual input not predicted by motor commands). Stimuli last for 1 second and are alternated with baseline phases (absence of visual input and motor predictions). **(c)** Left: Before plasticity, somatic excitation (light red) and inhibition (light blue) in PCs are not balanced. Excitatory and inhibitory currents shifted by ± 20 pA for visualization. The varying net excitatory current (gray) causes the PC population rate to deviate from baseline. Right: Response relative to baseline (Δ*R/R*) of all PCs in feedback (FB), mismatch (MM) and playback (PB) phase, sorted by amplitude of mismatch response. None of the PCs are classified as nPE neurons (indicated by gray shading to the right). **(d)** Same as in (c) after plasticity. Somatic excitation and inhibition are balanced. PC population rate remains at baseline. All PCs classified as nPE neurons (also indicated by black shading to the right).

### Negative prediction-error neurons emerge by balancing excitation and inhibition

Before the onset of plasticity, synaptic connections were randomly initialized, so PCs receive unbalanced excitation and inhibition. Therefore, all PCs change their firing rate in response to both feedback and playback stimuli, indicating the absence of nPE neurons (Fig. 1c). During quasi-natural sensorimotor experience, inhibitory plasticity strengthens or weakens inhibitory synapses to diminish the firing rate deviations of PCs from their baseline firing rate (Supplementary Fig. S1). At the same time, dendritic inhibition mediated by SOM interneurons was sufficiently strengthened to suppress the motor prediction arriving at the apical dendrite. After synaptic plasticity, somatic excitation and inhibition are balanced on a stimulus-by-stimulus basis (Fig. 1d). PCs merely show small and transient onset/offset responses to feedback and playback stimuli. In contrast, all PCs show an increase in activity for feedback mismatch stimuli (Fig. 1d). Hence, inhibitory synaptic plasticity generates nPE neurons by balancing excitation and inhibition in PCs for quasi-natural conditions.

### Balance of excitation, inhibition and disinhibition in different functional prediction circuits

It is not fully resolved which interneuron types receive sensory inputs, motor signals or both. The circuit we studied so far was motivated by the widely accepted view that PCs and SOM and PV interneurons show visual responses ^11,20–25^, while long-range (motor) predictions arrive in the superficial layers of V1 and target VIP neurons ^11,16,24,26^ and the apical and distal compartments of PCs ^11,23^. Because this view is not uncontested^26^, we systematically varied the inputs to the different neuron classes. We first studied circuit variations in which PCs and PV neurons receive visual and/or motor signals (Fig. 2, see also Supplementary Fig. S2).

**Figure 2.**
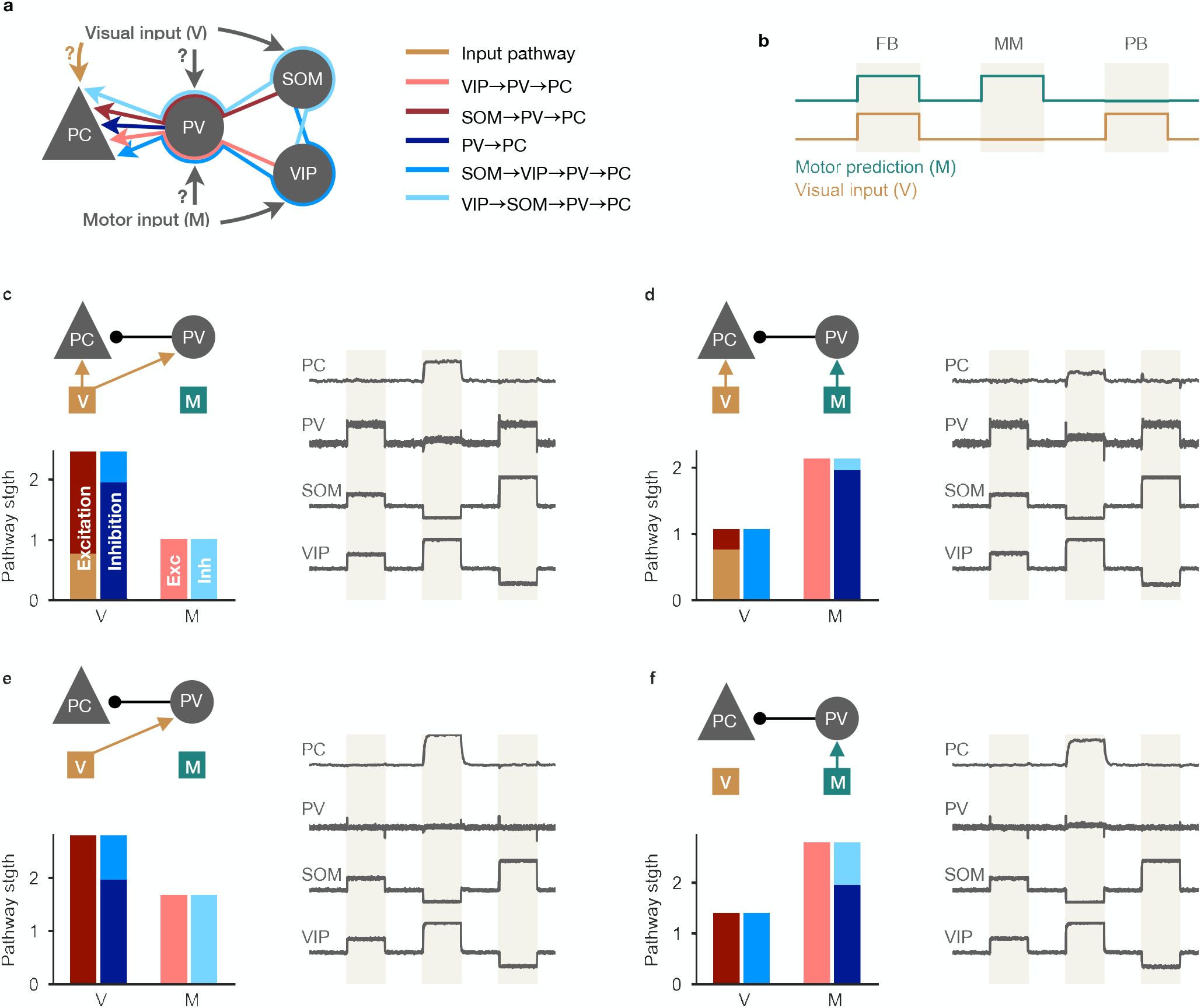
Multi-pathway balance of excitation and inhibition in different nPE neuron circuits. **(a)** Excitatory, inhibitory, disinhibitory and dis-disinhibitory pathways onto PCs that need to be balanced in nPE neuron circuits. Input to the soma of PCs and PV neurons is varied (c-f). SOM neurons receive visual input, VIP neurons receive a motor-related prediction. **(b)** Test stimuli: Feedback (FB), mismatch (MM) and playback (PB) phases of 1 second each. **(c)** PCs and PV neurons receive visual input (left, top). When all visual (V) and motor (M) pathways are balanced (left, bottom), PCs act as nPE neurons (right). PV neuron activity increases in both feedback and playback phases. Responses normalized between −1 and 1 such that baseline is zero. **(d)** Same as in (c) but PV neurons receive motor predictions. **(e)** Same as in (c) but PC s receive no visual input. PV neurons remain at baseline in the absence of visual input to the soma of PCs. **(f)** Same as in (c) but PCs receive no visual input and PV neurons receive motor predictions. PV neurons remain at baseline in the absence of visual input to the soma of PCs.

We found that inhibitory plasticity establishes nPE neurons independent of the input configuration onto PCs and PV neurons (Fig. 2b-e, right). The emerging connectivity of the interneuron circuits varied, however. For PCs not to respond above baseline in feedback and playback phase, various excitatory, inhibitory, disinhibitory and dis-disinhibitory pathways need to be balanced. An informative example is the input configuation in which PCs receive visual input and PV neurons receive motor predictions (Fig. 2c). In this case, visual inputs arrive at the PCs as direct excitation, as disinhibition through the SOM-PV pathway, and as dis-disinhibition via the SOM-VIP-PV pathway (Fig. 2a). To keep the PCs at their baseline during the playback phase, these three pathways need to be balanced (Fig. 2c, left). Similarly, motor signals arrive at the PCs as inhibition from PV neurons, dis-inhibition via the VIP-PV pathway, dis-dis-inhibition via the VIP-SOM-PV pathway and as direct excitation to the dendrite that is canceled by SOM-mediated inhibition. Again, all these pathways need to be balanced to keep the PCs at their baseline for fully predicted visual stimuli (Fig. 2c, left). Analog balancing arguments hold for other input configurations ((Fig. 2b-e, left).

While the flow of visual and motor information in the learned inhibitory microcircuit is different for different input configurations, the neural responses of the different interneuron classes provide limited information about the input configuration. PV neuron activity reflects whether PCs receive visual input: If PCs receive visual input, PV responses increase during feedback and playback phases to balance the sensory input at the soma of PCs (Fig. 2b-e, right). If PCs receive no visual input, PV neurons remain at their baseline firing rate (Fig. 2d-e, right). The activity of SOM and VIP neurons varies between playback, feedback and mismatch phases, but is independent of the input configuration for PCs and PV interneurons (Fig. 2b-e, right).

In summary, inhibitory plasticity can establish functional nPE circuits irrespective of the inputs onto the soma of PCs and PV neurons. Although the underlying circuits vary substantially in the specific balance of pathways, the neural activity patterns only weakly reflect the underlying information flow.

### Simulated optogenetic manipulations disambiguate prediction circuits

We hypothesized that the need to simultaneously balance several pathways offers a way to disambiguate the different prediction circuits by optogenetic manipulations. To test this, we systematically suppressed or activated PV, SOM and VIP interneurons in each input configuration after inhibitory plasticity had established the respective nPE circuit.

We found that in our model, such simulated optogenetic experiments are highly informative about the underlying input configuration (Fig. 3). For example, PV neuron inactivation changes the response of nPE neurons during feedback, playback and mismatch phases if and only if the PCs receive visual inputs. VIP inactivation renders nPE neurons silent unless PV neurons receive motor predictions, in which case they are transformed into positive prediction-error (pPE) neurons. Since SOM and VIP neurons are mutually inhibiting, the same information can be gained by an over-activation of SOM neurons that effectively silences VIP neurons.

**Figure 3.**
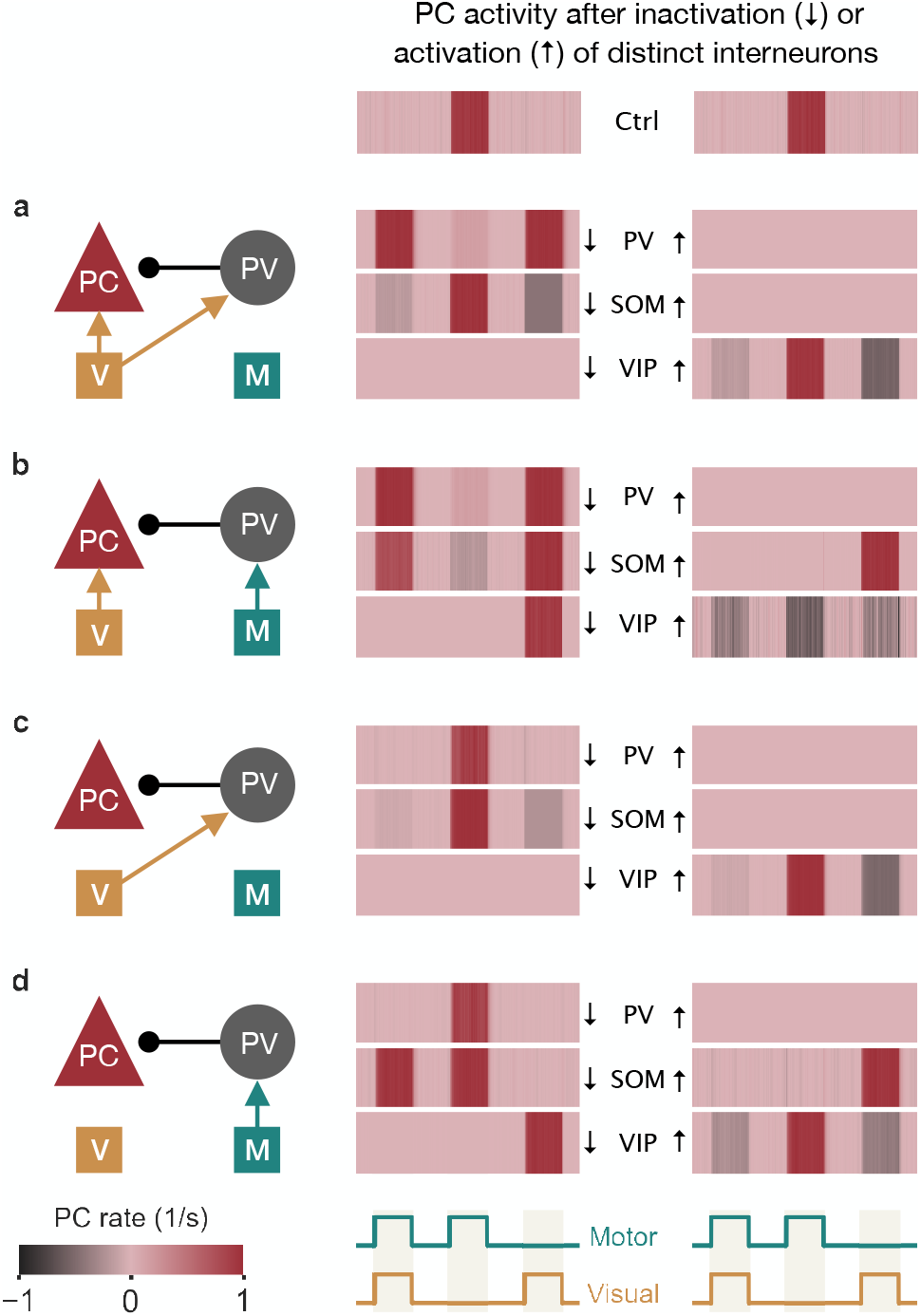
Simulated optogenetic manipulations of PV, SOM and VIP neurons disambiguate prediction-error circuits. **(a)** Left: nPE neuron circuit in which PCs and PV neurons receive visual input. Inactivation (middle) or activation (right) of PV (first row), SOM (second row) or VIP neurons (third row). Optogenetic manipulations change responses of nPE neurons (Ctrl) in feedback, mismatch and playback phases. Responses normalized between −1 and 1 such that baseline is zero. Inactivation input is −8*s*^−1^. Activation input is 5*s*^−1^. **(b)** Same as in (a) but PV neurons receive motor-related prediction. **(c)** Same as in (a) but PCs receive no visual input. **(d)** Same as in (a) but PCs receive no visual input and PV neurons receive a motor-related prediction.

In summary, our model predicts that optogenetic experiments unveil a unique fingerprint for nPE circuits that differ in their inputs onto PCs and PV neurons.

### Fraction of nPE neurons is modulated by inputs to SOM and VIP interneurons

In the model considered so far, all PCs developed into nPE neurons during learning, irrespective of the inputs to PCs and PV interneurons. However, nPE neurons represent only a small fraction of neurons in mouse V1 ^10,11^. Given that in our model, motor predictions arriving at the apical dendrites are canceled by SOM neuron-mediated inhibition, we hypothesized that the fraction of PCs that develop into nPE neurons depends on the distribution of visual and motor input onto SOM and VIP neurons.

To test this, we allow neurons of both SOM and VIP populations to receive either visual input or a motor prediction thereof. A fraction *f* of SOM neurons and a fraction (1 − *f*) of VIP neurons receive visual input. The remaining SOM and VIP neurons receive motor input (Fig. 4a). When the majority of SOM neurons receive visual inputs and the majority of VIP neurons receive motor predictions (*f* ≈ 1), all PCs develop into nPE neurons (Fig. 4b, left). Reducing the proportion of SOM neurons that receive visual input (and, equivalently, the proportion of VIP neurons that receive the motor prediction), the fraction of nPE neurons decreases (Fig. 4b, middle). Non-nPE neurons remain at their baseline in all three phases, show a suppression during mismatch or develop into pPE neurons that respond only during playback. pPE neurons only emerge when the inputs to SOM and VIP neurons are reversed such that most SOM neurons receive motor predictions (Fig. 4b, right).

**Figure 4.**
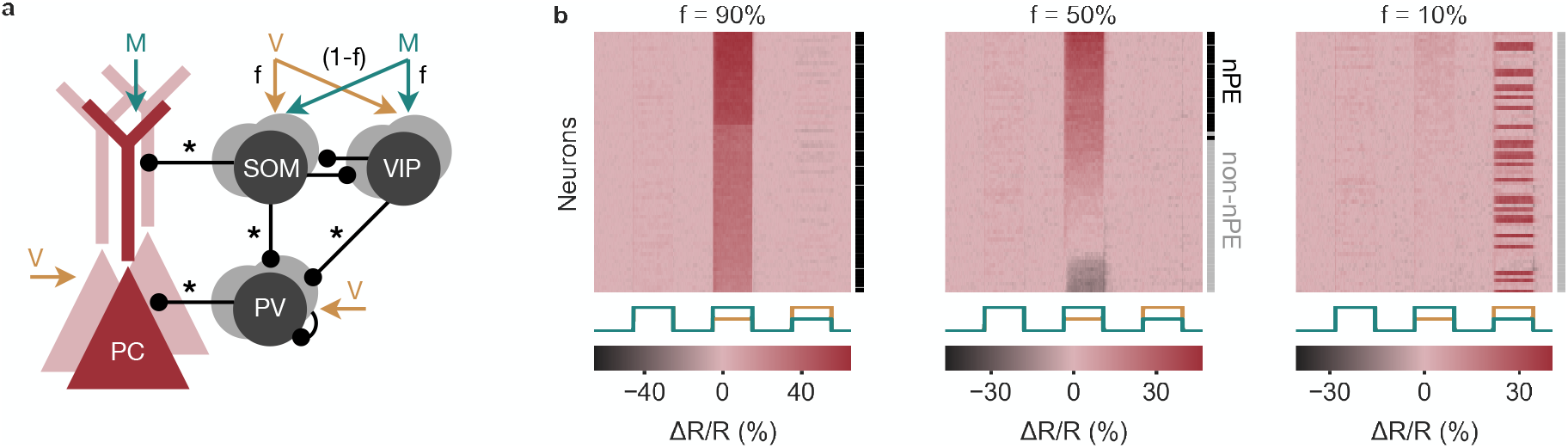
Fraction of nPE neurons depends on SOM and VIP neuron inputs. **(a)** Somatic compartment of PCs, PV neurons, a fraction *f* of SOM neurons and a fraction (1 − *f*) of VIP neurons receive visual input. The remaining SOM and VIP neurons receive motor predictions. **(b)** Response relative to baseline (Δ*R/R*) of all PCs in feedback, mismatch and playback phases, sorted by amplitude of mismatch response. The fraction of nPE neurons that develop during learning decreases with *f* (also indicated by black and gray shading to the right). The increasing fraction of non-nPE neurons comprises neurons that remain at their baseline in all three phases, show a suppression during mismatch or develop into positive prediction-error neurons that respond only during playback.

In summary, the fraction of nPE neurons that develop during learning depends on the distribution of visual input and motor predictions onto both SOM and VIP neurons.

### Experience-dependence of mismatch and interneuron responses

Attinger et al. ^11^ showed that the number of nPE neurons and the strength of their mismatch responses decrease when mice are trained in artificial conditions, in which motor predictions and visual flow were uncorrelated (“non-coupled training”). To test whether the model shows the same experience-dependence, we generated a modified training phase, in which visual inputs and motor-related predictions were statistically independent (Fig. 5a). We found that the number of nPE neurons and their mismatch responses also decrease for non-coupled trained relative to quasi-natural trained networks (Fig. 5b). This decrease is primarily due to changes in PCs and PV neurons, while the responses of SOM and VIP neurons during the mismatch phase are largely independent of the training paradigm (Fig. 5c). Hence, the experience-dependence of the model circuit is in line with that of nPE neurons in rodent V1 ^11^.

**Figure 5.**
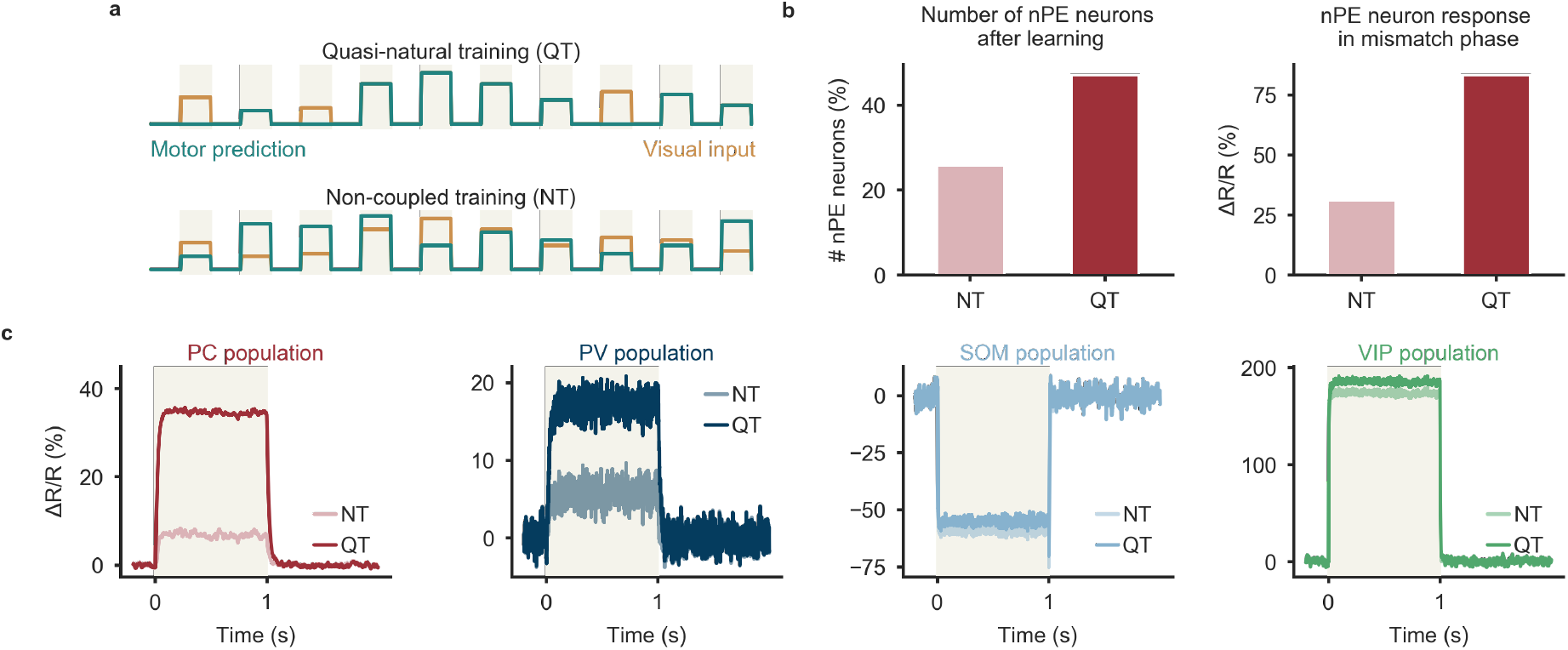
Experience-dependence of nPE and PV neurons. **(a)** The network is either exposed to a sequence of feedback and playback phases (quasi-natural training, QT) or to decoupled sensorimotor experience (non-coupled training, NT). **(b)** The number of nPE neurons that develop during learning (left) and their mismatch responses (right) are smaller for NT than for QT networks. **(c)** Population response (Δ*R/R*) of PCs, PV, SOM and VIP neurons during mismatch phase. SOM and VIP neurons show the same mismatch response for QT and NT, PCs and PV neurons show stronger responses in QT than in NT. Fraction of SOM neurons that receive visual input is f=80%.

### nPE circuits can also be learned by biologically plausible learning rules

In our model, nPE neurons developed though inhibitory plasticity that establishes an excitation-inhibition (E/I) balance in PCs. So far, we used learning rules that approximate a backpropagation of error ^27^, which changed SOM→PV and VIP→PV connections such as to minimize the difference between the PC firing rate and a baseline rate. The biological plausibility of such backpropagation rules, which are broadly used in artificial intelligence, is still debated, because they rely on information that is not locally available at the synapse in question ^28,29^. We therefore wondered whether prediction-error circuits can also be established by biologically plausible local learning rules.

We found that nPE neurons also emerged when the backpropagation rules were replaced by a form of plasticity that changes SOM→PV and VIP→PV synapses in proportion to the difference between the excitatory recurrent drive onto PV neurons and a target value (Fig. 6a). This local form of learning also balanced excitation and inhibition (Fig. 6b,c) and all PCs develop into nPE neurons (Fig. 6c).

**Figure 6.**
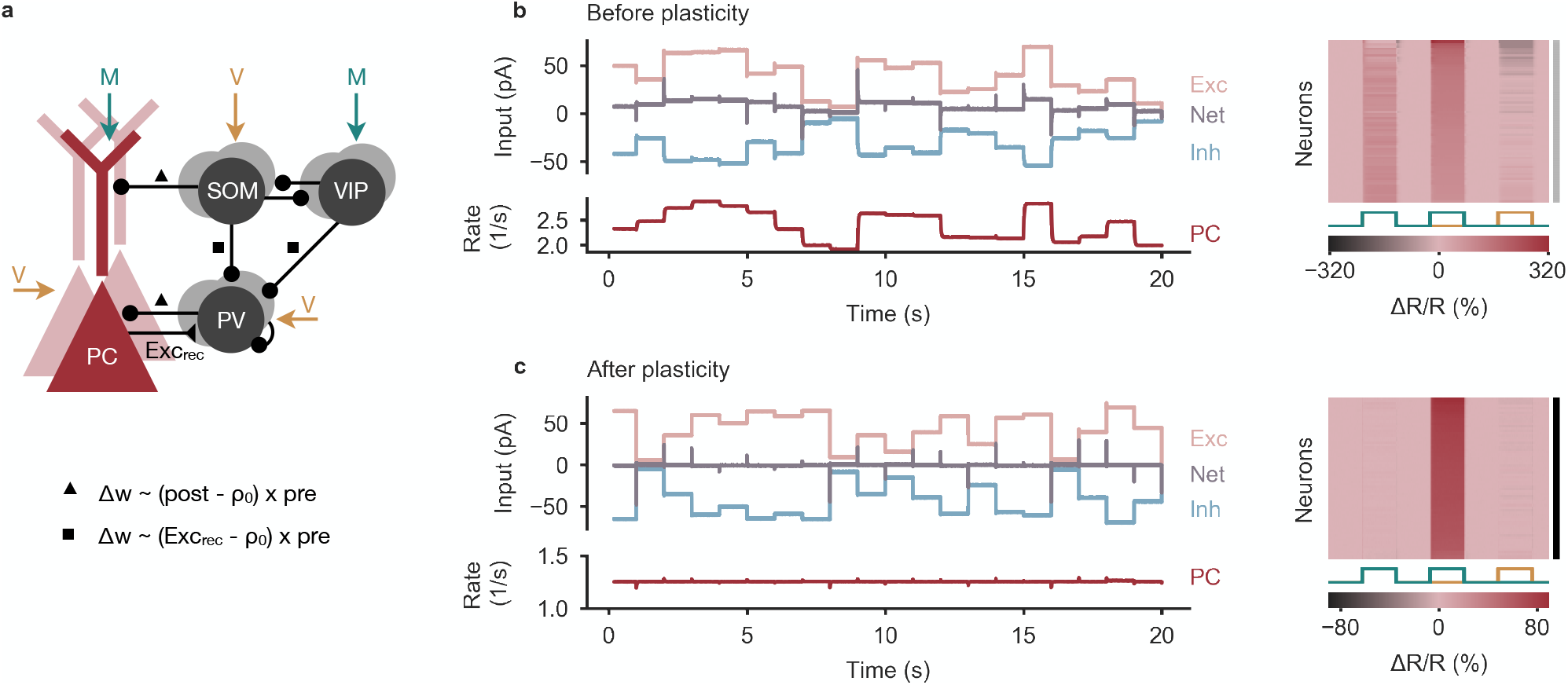
Learning nPE neurons by biologically plausible learning rules. **(a)** Network model as in Fig. 1. Connections marked with symbols undergo experience-dependent plasticity. Connections onto PCs follow inhibitory plasticity rule akin to Vogels et al. ^19^ (triangle). SOM→PV and VIP→PV synapses change in proportion to the difference between the excitatory recurrent drive onto PV neurons and a target value (square). **(b)** Left: Before plasticity1 somatic excitation (light red) and inhibition (light blue) in PCs are not balanced. Excitatory and inhibitory currents shifted by ±20 pA for visualization. The varying net excitatory current (gray) causes the PC population rate to deviate from baseline. Right: Response relative to baseline (Δ*R/R*) of all PCs in feedback1 mismatch and playback phases1 sorted by amplitude of mismatch response. None of the PCs are classified as nPE neurons (indicated by gray shading to the right). **(c)** Same as in (b) after plasticity. Somatic excitation and inhibition are balanced. PC population rate remains at baseline. All PCs classified as nPE neurons (also indicated by black shading to the right).

The plasticity rules can be further simplified when PCs do not receive visual information. In this case, the strength of SOM→PV and VIP→PV synapses can be learned according to a homeostatic rule 19 that aims to sustain a target rate in the PV neurons (Supplementary Fig. S3).

In summary, the backpropagation-like learning rules for the synapses onto PV neurons can be approximated by biologically plausible rules that exploit local information available at the respective synapses.

## Discussion

How the nervous system disentangles self-generated and external sensory stimuli is a long-standing question ^1,2,6^. Here1 we investigated the circuit level mechanisms that underlie the computation of prediction errors and how different types of inhibitory neurons shape these prediction circuits. We used computational modelling to show that nPE neurons can be learned by balancing excitation and inhibition in cortical microcircuits with three types of interneurons. We show that the required E/I balance can be achieved by biologically plausible forms of synaptic plasticity. Furthermore1 the experience-dependence of the circuit is similar to that of nPE circuits in mouse V1 ^11^.

Our model makes a number of predictions. Firstly, the multi-pathway balance of excitation and inhibition suggests that the input configuration of the prediction circuit could be disambiguated using cell type-specific modulations of neural activity. This could be achieved by optogenetic or pharmacogenetic manipulations, or by exploiting the differential sensitivity of interneuron classes to neuromodulators. The precarious nature of an exact multi-pathway balance also suggests that nPE neurons might change their response characteristics in a context-dependent way, e.g., by neuromodulatory effects.

Secondly, the central assumption of the model is that nPE neurons emerge by a self-organized E/I balance during sensorimotor experience. It therefore predicts that (i) sensorimotor experience that the animal is habituated to should lead to balanced excitation and inhibition in PCs, (ii) E/I balance should break for sensorimotor experience the animal has rarely encountered, e.g., for mismatches of sensory stimuli and motor predictions and (iii) during altered sensorimotor experience in a virtual reality setting or when the excitability of specific interneuron types is altered, interneuron circuits should gradually reconfigure to reestablish the E/I balance.

During learning, we exposed the network to sensory inputs and motor-related predictions designed to reflect coupled sensorimotor experience. To allow for changes in the external world that do not arise from the animal’s own movements, we included “playback” phases in which the visual input is stronger than predicted by the motor-related input. Consistent with the experimental setup of Attinger et al. ^11^, we deliberately excluded feedback mismatch phases. In the model, the stimuli experienced during learning have a strong impact on the response structure of the PCs, because the learning rules aim to keep the PCs at a given baseline rate at all times. The inclusion of feedback and playback phases during learning therefore leads to neurons that remain at their baseline during those phases, in line with nPE neurons. In mouse V1, nPE neurons exhibit an average rate decrease during playback when the animals were only exposed to perfectly coupled sensorimotor experience ^11^. When our network was trained in the same way, we also observed that PCs reduced their firing rate during playback phases (Supplementary Fig. S4). This can be a result of an excess of somatic inhibition, dendritic inhibition or both. The model hence predicts that the rate reduction during playback phases observed by Attinger et al. ^11^ vanishes when playback phases are included during training.

The interneuron circuit in our model is motivated by the canonical circuit found in a variety of brain regions ^17,18,30^. In addition to the connections between interneuron classes that are frequently reported as strong and numerous, we included VIP→PV synapses in the circuit, because a mathematical analysis reveals that they are required for a perfect E/I balance during both feedback and playback phases (see Supplementary Notes). While VIP→PV synapses have been found in visual ^17^, auditory ^31^, somatosensory ^30,32^ and medial prefrontal cortex ^31^, as well as amygdala ^33^, they are less prominent and often weaker than SOM→PV connections (but see Krabbe et al. ^33^). VIP→PV synapses can be excluded when the conditions for nPE neurons during feedback and playback phases are mildly relaxed^10,11,13^ and when PV neurons receive visual, but not motor inputs (Supplementary Fig. S5).

We used a mathematical analysis to identify a number of synapses in the circuit that undergo experience-dependent changes. While the synapses from PV neurons onto PCs established a baseline firing rate in the absence of visual input and motor predictions, the synergy between the SOM→PV, VIP→PV and SOM→PC synapses guaranteed that the baseline is retained in feedback and playback phase. Our mathematical analysis unveiled constraints for the interneuron motif, that is, the relation between the strengths of a number of inhibitory synapses (see Methods, Eqs. 8, 9). The multi-pathway balance of excitation and inhibition could also be achieved by synaptic plasticity in other inhibitory synapses – for example the mutual inhibition between SOM and VIP neurons. However, the assumption that mainly the inhibitory synapses onto PV neurons are plastic is supported by the observation that PV neuron activity – in contrast to SOM and VIP neuron activity – is experience-dependent^11^.

In the model, the plastic inhibitory synapses onto PV neurons change according to non-local information that might not be directly available at the synapse. These synapses therefore implement an approximation of a backpropagation of error, the biological plausibility of which is debated ^28^. We showed that this plasticity rule can be approximated by biologically plausible variants of the plasticity rules. If PCs do not receive direct visual input (Supplementary Fig. S3), the backpropagation-like algorithm can be replaced by a simple homeostatic Hebbian plasticity rule in the synapses onto the PV interneurons. Given that PCs in V1 are known to receive substantial visual drive ^21,22^, this assumption is unlikely to be valid. We therefore propose an alternative form of plasticity that changes SOM→PV and VIP→PV synapses in proportion to the difference between the excitatory recurrent drive onto PV neurons and a target value (Fig. 6). The underlying mechanism is similar to feedback alignment^34^ and requires sufficient overlap between the set of postsynaptic PCs a PV neuron inhibits and the set of presynaptic PCs the same PV neuron receives excitation from. This is likely, given the high connection probability between PCs and PV neurons ^17,18,35^.

We modelled the apical dendrite of PCs as a single compartment that integrates excitatory and inhibitory input currents and has the potential to produce calcium spike-like events^36–39^. Moreover, we assumed that an overshoot of inhibition decouples the apical tuft of the PCs from their soma, by including a rectifying non-linearity that precludes an excess of dendritic inhibition to influence somatic activity. However, the presence or nature of these dendritic nonlinearities has a minor influence on the development of nPE neurons (Supplementary Fig. S6). When we allowed dendritic inhibition to influence the soma, inhibitory plasticity still established nPE neurons, although the learned interneuron circuit differs with respect to the synaptic strengths. The additional dendritic inhibition reduces the required amount of somatic, PV-mediated inhibition. This is primarily the case during playback phases, when the excitatory motor input to the apical dendrite is absent. PV neurons are therefore less active during the playback phase than during the feedback phase (Supplementary Fig. S6), consistent with recordings in mouse V1 ^11^.

By modelling the apical dendrite as a single compartment, we also neglected the possibility that dendritic branches process distinct information. However, we expect that the suggested framework of generating predictive signals by a compartment-specific E/I balance generalizes to more complex dendritic configurations, in which local inhibition could contribute by gating different dendritic inputs^40^.

Cortical circuits are complex and contain a large variety of interneuron classes ^15,16,18^. We restricted the model to three of these classes: PV, SOM and VIP neurons. It is conceivable that several other interneuron types can play a pivotal role in prediction-error circuits. The dendrites of layer 2/3 neurons reach out to layer 1, the major target for feedback connections ^23,41,42^ and home to a number of distinct interneuron types ^43,44^, which may contribute to associative learning ^45,46^. In particular, NDNF neurons unspecifically inhibit apical dendrites located in the superficial layers, and at the same time receive strong inhibition from SOM neurons ^45^. Hence, it is possible that these interneurons also shape the processing of feedback information, including the computation of prediction errors.

PCs in L2/3 of V1 have very low spontaneous firing rates^22,47^. A potential rate decrease during feedback and playback could hence be hard to detect. Whether the low response of nPE neurons during feedback and playback phases are due to an E/I balance – as suggested here – or due to an excess of inhibition may hence be difficult to decide, and could for example be resolved by intracellular recordings.

Our model suggests a well-orchestrated division of labor of PV, SOM and VIP interneurons that is shaped by experience: While PV neurons balance the sensory input at the somatic compartment of PCs, SOM neurons cancel feedback signals at the apical dendrites. VIP neurons ensure sufficiently large mismatch responses by amplifying small differences between feedforward and feedback inputs^11,39^. Given the relative uniformity of cortex in its appearance, structure and cell types ^48,49^, it is conceivable that the same principles also hold for other regions of the cortex beyond V1. Shedding light on the mechanisms that constitute the predictive power of neuronal circuits may in the long run contribute to an understanding of psychiatric disorders that have long been associated with a malfunction of the brain’s prediction machinery ^50–52^ and specific types of interneurons ^53–55^.

## Methods

### Network model

We simulated a rate-based network model of excitatory pyramidal cells (*N*_PC_ = 70) and inhibitory PV, SOM and VIP neurons (*N*_PV_ = *N*_SOM_ = *N*_VIP_ = 10). All neurons are randomly connected with connection strengths and probabilities given below (see “Connectivity”).

The excitatory pyramidal cells are described by a two-compartment rate model that was introduced by Murayama et al. ^38^. The dynamics of the firing rate *r*_E,*i*_ of the somatic compartment of neuron *i* obeys

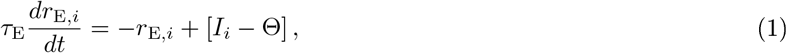

where *τ*_E_ denotes the excitatory rate time constant (*τ*_E_=60 ms), Θ terms the rheobase of the neuron (Θ = 14 *s*^−1^). Firing rates are rectified to ensure positivity. *I*_*i*_ is the total somatic input generated by somatic and dendritic synaptic events and potential dendritic calcium spikes:

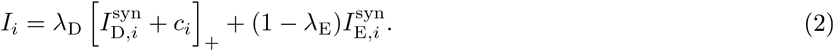

Here, the function [*x*]_+_ = max(*x*, 0) is a rectifying nonlinearity that prohibits an excess of inhibition at the apical dendrite to reach the soma. 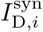 and 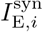 are the total synaptic inputs into dendrite and soma, respectively, and *c*_*i*_ denotes a dendritic calcium event. *λ*_D_ and *λ*_E_ are the fraction of “currents” leaking away from dendrites and soma, respectively (*λ*_D_=0.27, *λ*_E_=0.31). The synaptic input to the soma 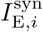 is given by the sum of external sensory inputs *x*_E_ and PV neuron-induced (P) inhibition,

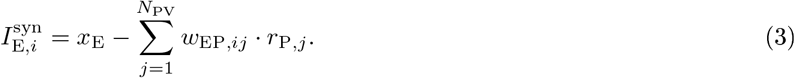

The dendritic input 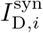 is the sum of motor-related predictions *x*_D_, the recurrent, excitatory connections from other PCs and SOM neuron-induced (S) inhibition:

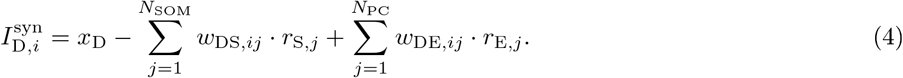

The weight matrices *w*_EP_, *w*_DS_ and *w*_DE_ denote the strength of connection between PV neurons and the soma of PCs (*w*_EP_), SOM neurons and the dendrites of PCs (*w*_DS_) and the recurrence between PCs (*w*_DE_), respectively. The input generated by a calcium spike is given by

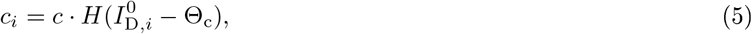

where *c* scales the amount of current produced (*c* = 7 *s*^−1^), *H* is the Heaviside step function, Θ_c_ represents a threshold that describes the minimal input needed to produce a Ca^2+^-spike (Θ_c_ = 28 *s*^−1^) and 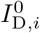 denotes the total, synaptically generated input in the dendrites,

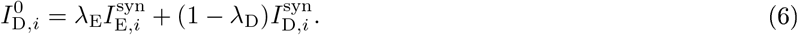

Note that we incorporated the gain factor present in Murayama et al. ^38^ into the parameters to achieve unit consistency for all neuron types.

The firing rate dynamics of each interneuron is modeled by a rectified, linear differential equation ^56^,

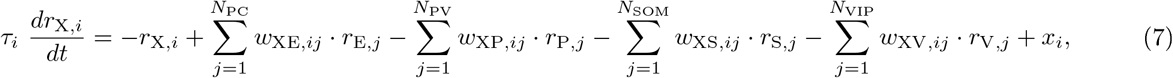

where *r*_X,*i*_ denotes the firing rate of neuron *i* from neuron type *X* (*X* ∈ {*P, S, V*}) and *x*_*i*_ represents external inputs. The weight matrices *w*_XY_ denote the strength of connection between the presynaptic neuron population *Y* and the postsynaptic neuron population *X*. The rate time constant *τ*_*i*_ was chosen to resemble a fast GABA_A_ time constant, and set to 2 ms for all interneuron types included.

### Negative prediction-error neurons

We define PCs as nPE neurons when they exclusively increase their firing rate during feedback mismatch (visual input smaller than predicted), while remaining at their baseline during feedback and playback phases. In a linearized, homogeneous network and under the assumption that the apical dendrites are sufficiently inhibited during feedback and playback phase, this definition is equivalent to two constraints on the interneuron network (see Supporting Information for a detailed analysis and derivation):

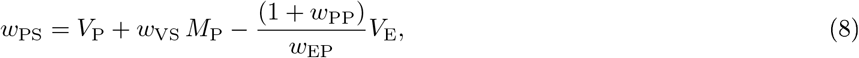

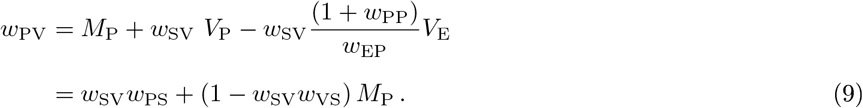

The parameters *V*_*X*_, *M*_*X*_ ∈ {0, 1} indicate whether neuron type *X* receives visual and motor-related inputs, respectively, and control the different input configurations. In addition to the conditions Eqs. 8 and 9, the synapses from SOM neurons onto the apical dendrites must be sufficiently strong to cancel potential excitatory inputs during feedback and playback phase.

In practice, we classify PCs as nPE neurons when Δ*R/R* is larger than 20% in the mismatch phase and less than ±10% elsewhere (Δ*R/R* = (*r* − *r*_BL_)*/r*_BL_, *r*_BL_: baseline firing rate). Tolerating small deviations in feedback and playback phase is more in line with experimental approaches. The results do not rely on the precise thresholds used for the classification.

### Connectivity

All neurons are randomly connected with connection probabilities motivated by the experimental literature^17,18,30,31,35,57–59^,

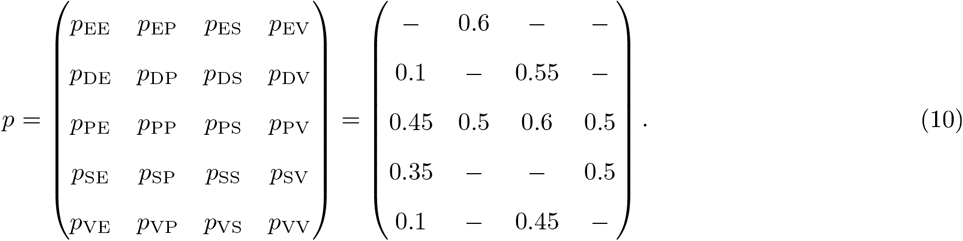

All cells of the same neuron type have the same number of incoming connections. The mean connection strengths are given by

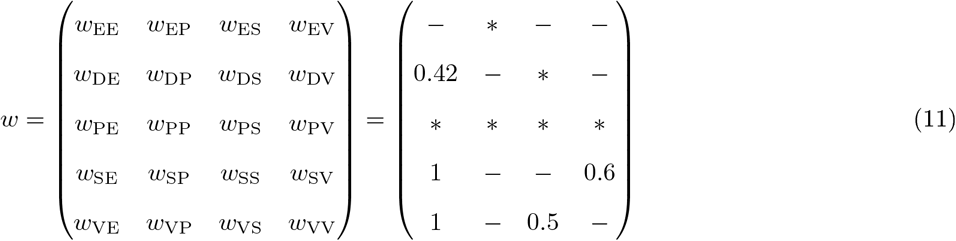

where the symbol * denotes weights that vary between simulations (e.g., subject to plasticity or computed from the equations (8) and (9)). For non-plastic networks, these synaptic strengths are given by *w*_EP_ = 2.8, *w*_DS_ = 3.5, *w*_PE_ = 1.5, *w*_PP_ = 0.1 (if PCs receive visual input) or *w*_PP_ = 1.5 (if PCs receive no visual input), *w*_PS_ and *w*_PV_ are computed from the equations (8) and (9).

For plastic networks, the initial connections between neurons are drawn from uniform distributions 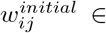 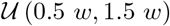 where *w* denotes the mean connection strengths given in (11) and *w*_EP_ = 1.75, *w*_EP_ = 0.35, *w*_PE_ = 2.5 (if PCs receive visual input) or *w*_PE_ = 1.2 (if PCs receive no visual input), *w*_PP_ = 0.5 (if PCs receive visual input) or *w*_PP_ = 1.5 (if PCs receive no visual input), *w*_PS_ = 0.3 and *w*_PV_ = 0.6. Please note that the system is robust to the choice of connections strengths. The connection strengths are merely chosen such that the solutions of Eqs. 8 and 9 comply with Dale’s principle.

All weights are scaled in proportion to the number of existing connections (i.e., the product of the number of presynaptic neurons and the connection probability), so that the results are independent of the population size.

### Inputs

All neurons receive constant, external background input that ensures reasonable baseline firing rates in the absence of visual and motor-related input. In the case of non-plastic networks, these inputs were set such that the baseline firing rates are *r*_E_ = 1*s*^−1^, *r*_P_ = 2*s*^−1^, *r*_S_ = 2*s*^−1^ and *r*_V_ = 4*s*^−1^. In the case of plastic networks, we set the external inputs to *x*_E_ = 28*s*^−1^, *x*_D_ = 0*s*^−1^, *x*_P_ = 2*s*^−1^, *x*_S_ = 2*s*^−1^ and *x*_V_ = 2*s*^−1^ (if not stated otherwise). In addition to the external background inputs, the neurons receive either visual input (*v*), a motor-related prediction thereof (*m*) or both.

In line with the experimental setup of Attinger et al. ^11^, we distinguish between baseline (*m* = *v* = 0), feedback (*m* = *v* > 0), feedback mismatch (*m* > *v*) and playback (*m* < *v*) phases. During training, the network is exposed to feedback and playback phases with stimuli drawn from a uniform distribution from the interval [0, 7*s*^−1^]. After learning, the strength of stimuli is set to 7*s*^−1^ (plastic networks) or 3.5*s*^−1^ (non-plastic networks).

### Plasticity

In plastic networks, a number of connections between neurons are subject to experience-dependent changes in order to establish an E/I balance for PCs. PV→PC and the PC→PV synapses establish the target firing rates for PCs and PV neurons, respectively. VIP→PV and SOM→PV synapses and the synapses from SOM neurons onto the apical dendrites of PCs ensure that PCs remain at their baseline during feedback and playback phase. The corresponding plasticity rules are of the form

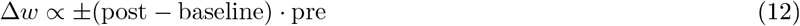

In detail, the connections from PV and SOM neurons onto the soma and the apical dendrites, respectively, obey inhibitory Hebbian plasticity rules akin to Vogels et al. ^19^

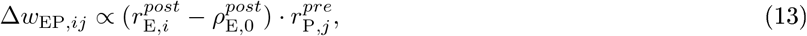

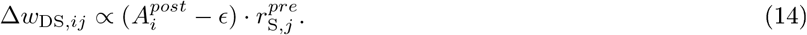

The parameter 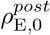 denotes the baseline firing rate of the postsynaptic PC, and the dendritic activity 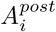 is given by the rectified synaptic events at the dendrites

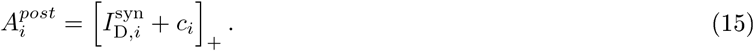

The small “correction” term *ε* eases the effect of strong onset responses (here, we used *ϵ* = 0.1*s*^−1^).

The connections from both SOM and VIP neurons onto PV neurons implement an approximation of a backpropagation of error

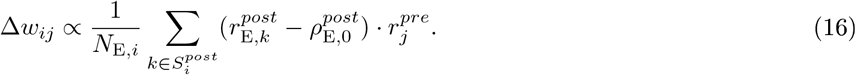

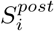 denotes the set of postsynaptic PCs a particular PV neuron is connected to, and *N*_E,*i*_ is the number of excitatory neurons in 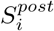.

When the connection probability between PCs and PV neurons is large, this backpropagation of error can be replaced by a biologically plausible learning rule that only relies on local information available in the PV neurons.

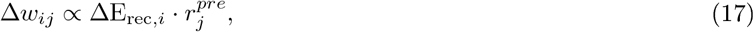

where ΔE_rec,*i*_ denotes the difference between the excitatory recurrent drive onto PV neuron *i* and a target value

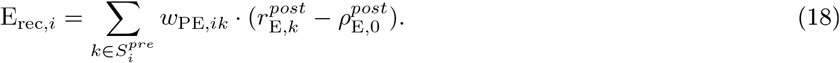

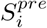 denotes the set of presynaptic PCs a particular PV neuron receives excitation from.

When nPE neurons do not receive direct visual input, the backpropagation rules can be simplified even further. The synapses onto PV neurons can be learned according to a Hebbian inhibitory plasticity rule ^19^ that aims to sustain a baseline rate in the PV neurons

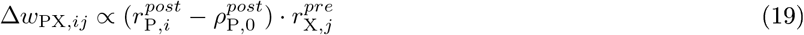

with *X* ∈ {*S*, *V*}. This baseline rate is established by modifying the connections from PCs onto PV neurons according to an anti-Hebbian plasticity rule

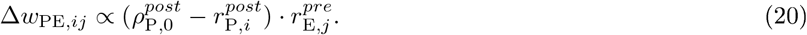

### Simulation

All simulations were performed in customized Python code written by LH. Differential equations were numerically integrated using a 2^nd^-order Runge-Kutta method with time steps between 0.05 and 2 ms. Neurons were initialized with *r*_*i*_(0) = 0. Source code will be made publicly available upon publication.

## Acknowledgements

We are grateful to Laura Bella Naumann and Joram Keijser for critical reading of the manuscript. The project is funded by the German Federal Ministry for Education and Research, FKZ 01GQ1201 and the DFG via the collaborative research center FOR 2143.

## Author Contributions

L.H. and H.S. conceived the project and designed the experiments. L.H. performed the simulations and mathematical analyses. L.H and H.S. interpreted the results and wrote the paper.

## Supplementary Information

### Supplementary Notes

We performed a mathematical analysis of a simplified model to identify the constraints that are imposed on the interneuron circuit by the presence of nPE neurons. We first describe the assumptions made and the definition of nPE neurons. We then derive the constraints for a simplified network with canonical interneuron connectivity including VIP-to-PV synapses. The solutions provide the relationship for the strength of synapses between different neuron types that must be satisfied for nPE neurons to emerge. We then show that the same network without VIP-to-PV synapses can only produce nPE neurons under very restrictive assumptions.

#### Constraints for the interneuron circuit

To derive the constraints for the interneuron network that are imposed by the presence of nPE neurons, we performed a mathematical analysis of a simplified network model, in which the nonlinearity of the dendritic compartment and the rectifying nonlinearities are neglected. This reduces the network to an analytically tractable linear system. The simplifications rely on the following assumptions:

1. During baseline, feedback and playback phases, SOM interneuron-mediated inhibition exceeds excitatory motor predictions arriving at the apical dendrites of PCs.
2. Any excess of inhibition in the dendrite does not affect the the soma of PCs.
3. During baseline, feedback and playback phases, all neuron types have positive firing rates, such that the rate rectification can be neglected.

These assumptions allow us to omit the dendritic compartment of PCs and consequently all synapses thereto. The remaining system of linear equations describes the activity of all neuron types during baseline, feedback and playback phase. For the subsequent analysis, we furthermore consider a homogeneous network, that is, all weights, neuronal properties and the number of incoming connections for cells of the same type are the same. As a result, we can reduce the high-dimensional system to 4 equations, each describing the dynamics of one representative firing rate per neuron type:

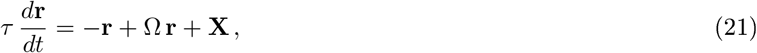

where *τ* denotes the rate time constant, **r** = [*r*_E_, *r*_P_, *r*_S_, *r*_V_]^*T*^ (subscripts refer to the different neuron types; E: soma of PC, P: PV, S: SOM, V: VIP), Ω is the weight matrix and **X** denotes the external inputs. In the steady state, the firing rates are given by

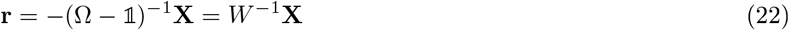

with the effective connectivity matrix *W* that includes the leak:

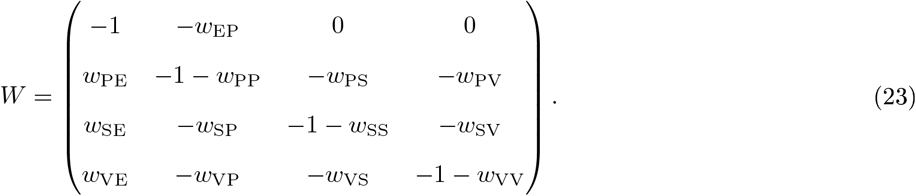

The weight parameters *w*_*XY*_ between neuron types are strictly positive to maintain the excitatory/inhibitory nature of the various neuron types.

In our model, an excitatory neuron is classified as a perfect nPE neuron, if

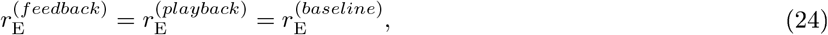

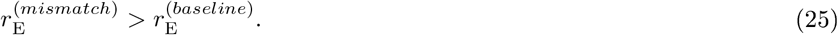

During feedback mismatch, the PC firing rate increases with respect to the baseline as long as the motor-related excitatory inputs exceed the somatic inhibition mediated by PV neurons. The conditions according to which no change in activity occurs in either feedback or playback phase (see Eq. 24) impose constraints on the weight configuration that need to be satisfied. These can be summarized by

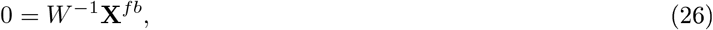

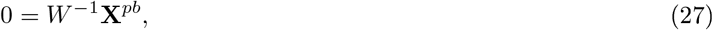

where **X**^*fb*^ and **X**^*pb*^ denote the excess external inputs above baseline during feedback and playback phase, respectively,

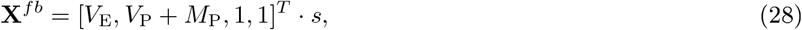

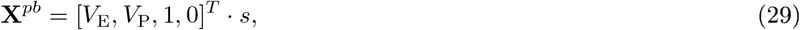

with *s* representing a varying excitatory stimulus strength. The parameters *V*_*X*_, *M*_*X*_ ∈ {0, 1} indicate whether neuron type *X* receives visual and motor-related inputs, respectively, and control the different input configurations.

##### Canonical interneuron connectivity with VIP-to-PV synapses

We start with the connectivity motif proposed by Pfeffer et al. ^17^. We also allow for connections from VIP to PV neurons. Although they are considered to be less prominent and weaker than connections from VIP to SOM neurons and are therefore often neglected in diagrams and computational models, those synapses have been observed in various brain regions ^17,30–33^. To this end, the respective connectivity matrix is given by

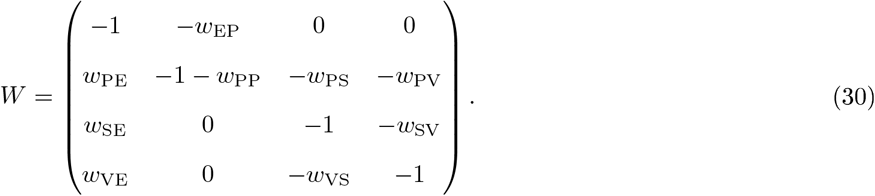

The constraints (26) and (27) defining nPE neurons are then given by

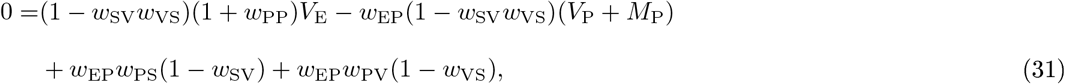

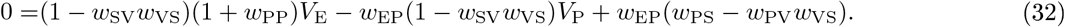

These two equations yield

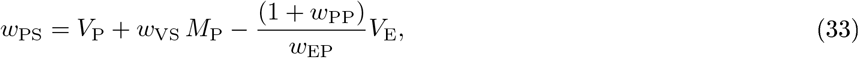

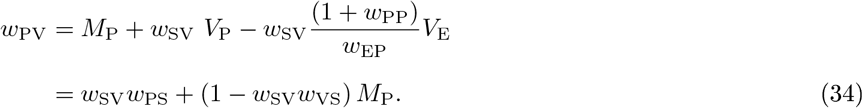

Eq. 33 and 34 are the mathematical formulation of the E/I balance of multiple pathways shown in Fig. 2 and Supplementary Fig. S2.

For the derivation above, we have assumed that the motor-related input is switched off during the playback phase. This assumption, however, can be relaxed. When motor predictions are merely smaller than the actual sensory input but non-zero during playback, analogous calculations yield the same constraints.

##### Canonical interneuron connectivity without VIP-to-PV synapses

Without connections from VIP onto PV neurons, the constraints (26) and (27) yield

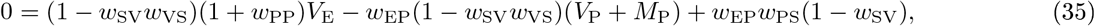

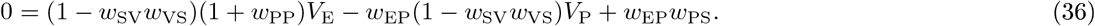

These two equations simplify to

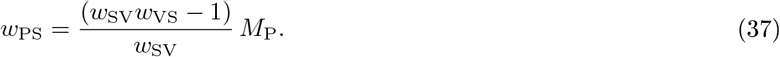

As the weight *w*_*PS*_ is strictly positive (see definition of weight matrix above), the product *w*_SV_*w*_VS_ must be larger than 1. This, however, indicates that networks with rate rectification exceed a bifurcation point and run into a winner-take-all (WTA) regime, in which either VIP or SOM neurons are silent^39^.

With VIP neurons being silent in all phases but during feedback mismatch phases, the constraint on *w*_PS_ can be recalculated from Eqs. 22 and 24 while neglecting VIP neurons:

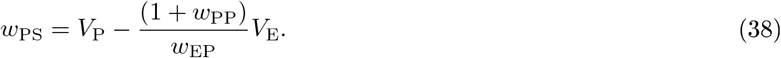

This equation reveals that PV neurons must receive visual input to ensure *w*_PS_ > 0.

In summary, this mathematical analysis shows that perfect nPE neurons can only emerge when VIP neurons are silent during all phases but the feedback mismatch phase.

Please note that the same results are obtained even if connections from PV to both SOM and VIP neurons are included.

## Supplementary Figures

**Figure S1.**
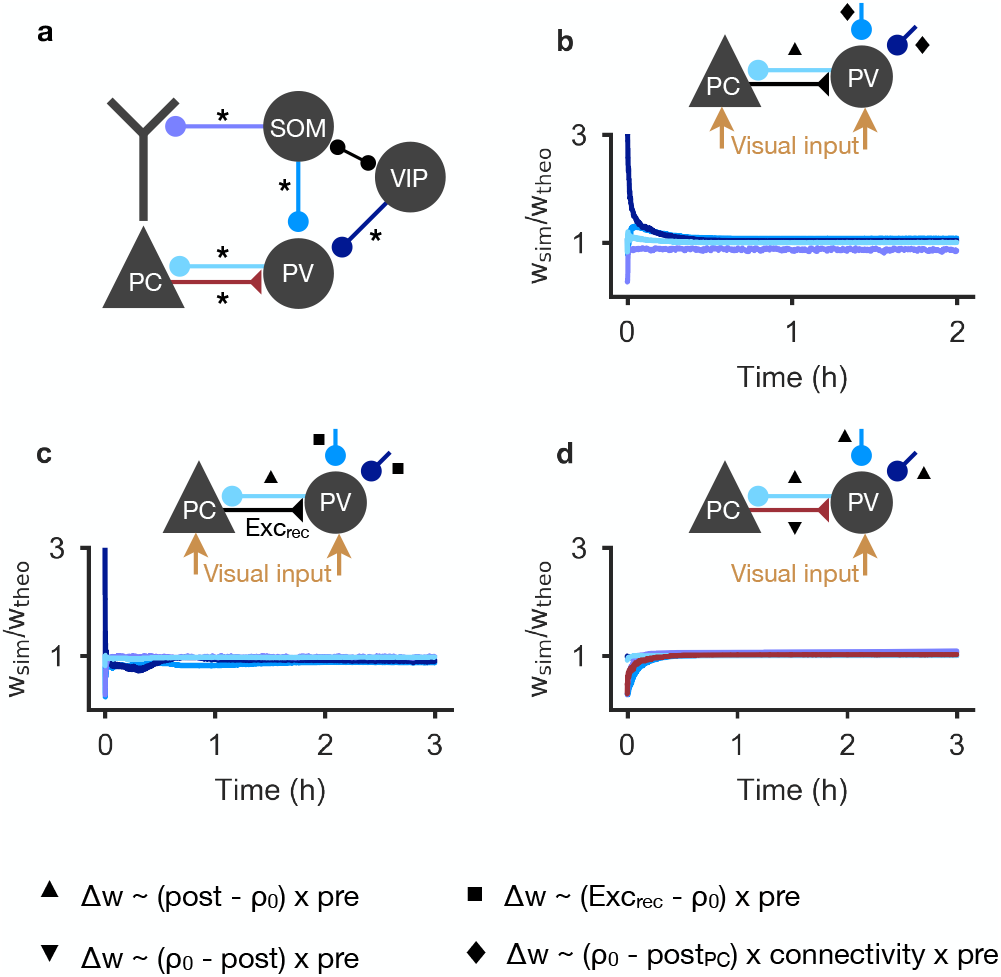
Learning of prediction-error circuits with different forms of homeostatic plasticity. **(a)** Network model as in Fig. 1. Connections colored and marked with an asterisk undergo experience-dependent plasticity. **b** PCs receive visual input. Connections onto PCs follow an inhibitory plasticity rule akin to Vogels et al. ^19^ (triangle). SOM→PV and VIP→PV synapses approximate a back-propagation of error (diamond). The averaged weights converge to a steady-state. Weights are normalized to the theoretically derived values for nPE neurons (see Methods). **(c)** Same as in (b) but SOM→PV and VIP→PV synapses change in proportion to the difference between the excitatory recurrent drive onto PV neurons and a target value (square). **(d)** Same as in (b) but visual drive onto PCs is absent. SOM→PV and VIP→PV synapses follow an inhibitory plasticity rule akin to Vogels et al. ^19^ (triangle). Connections from PCs onto PV neurons establish a baseline for PV neurons by an anti-Hebbian plasticity rule (inverted triangle).

**Figure S2.**
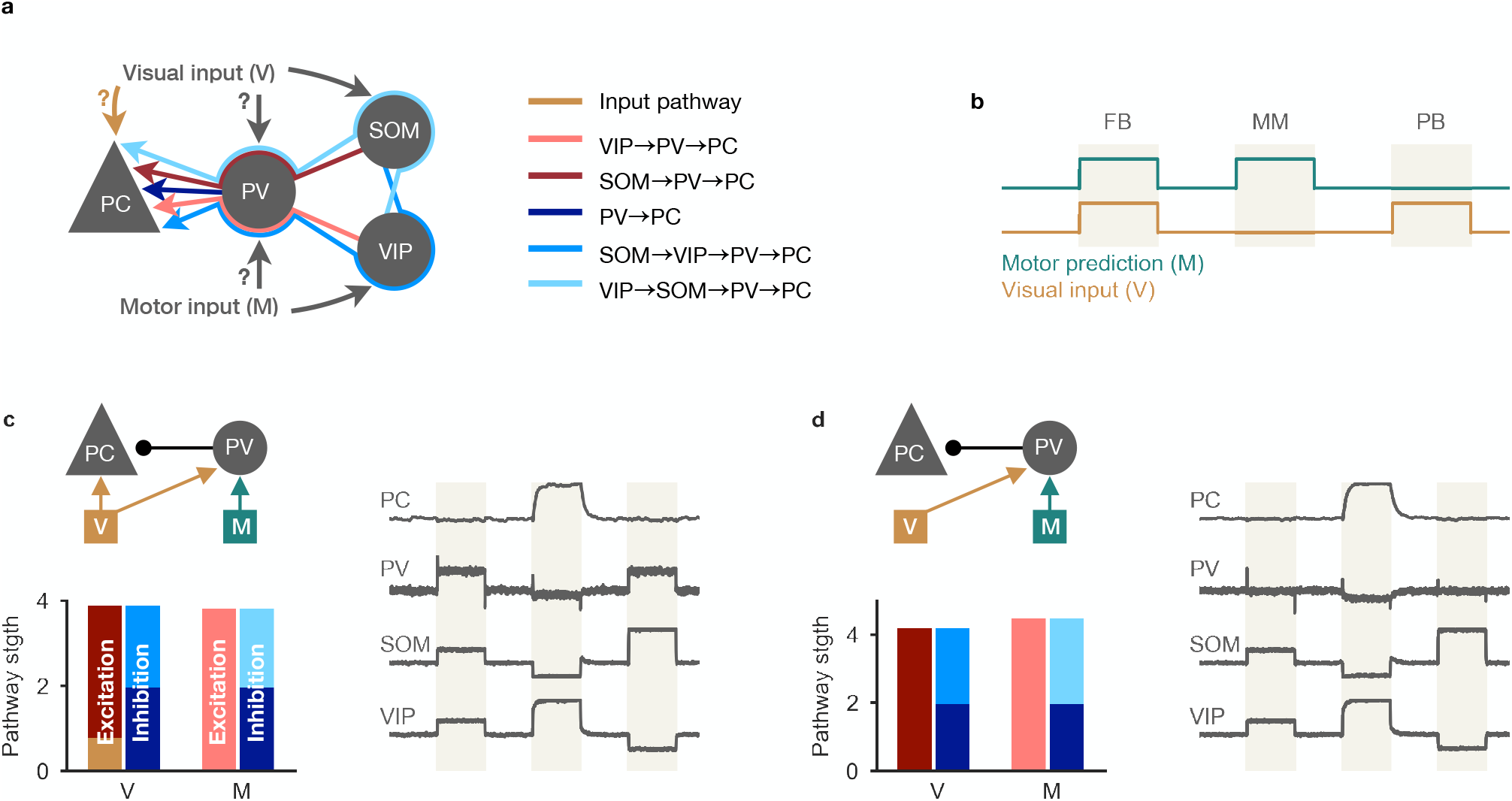
Multi-pathway balance of excitation and inhibition in different nPE neuron circuits with visual and motor input onto PV neurons. **a** Excitatory, inhibitory, disinhibitory and dis-disinhibitory pathways onto PCs that need to be balanced in nPE neuron circuits. Input to the soma of PCs and PV neurons is varied (b-c). SOM neurons receive visual input, VIP neurons receive a motor-related prediction. **(b)** Test stimuli: Feedback (FB), mismatch (MM) and playback (PB) phases of 1 second each. **c** PCs receive visual input (left, top). When all visual (V) and motor (M) pathways are balanced (left, bottom), PCs act as nPE neurons (right). PV neuron activity increases in both feedback and playback phases. Responses normalized between −1 and 1 such that baseline is zero. **d** Same as in (c) but PC receive no visual input. PV neurons remain at baseline in the absence of visual input to the soma of PCs.

**Figure S3.**
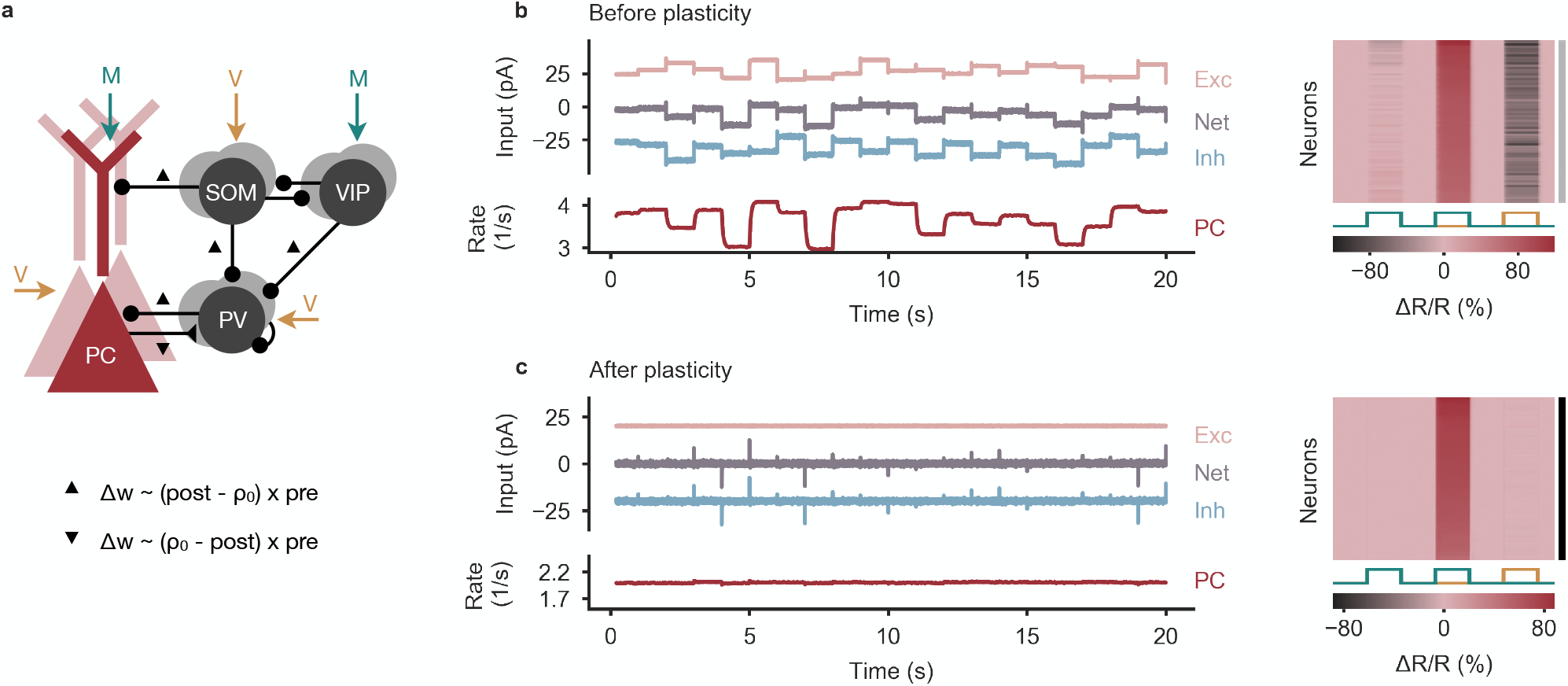
Learning nPE neurons by biologically plausible learning rules in networks without visual input at the soma of PCs. **a** Network model as in Fig. 1. Connections marked with symbols undergo experience-dependent plasticity. Inhibitory connections onto PCs and PV neurons follow inhibitory plasticity rule akin to Vogels et al. ^19^ (triangle). Synapses from PCs onto PV neurons follow an anti-Hebbian plasticity rule (inverted triangle). **b** Left: Before plasticity, somatic excitation (light red) and inhibition (light blue) at PCs are not balanced. Excitatory and inhibitory currents are shifted by ± 20 pA for visualization. The varying net excitatory current (gray) causes the PC population rate to deviate from baseline. Right: Response relative to baseline (Δ*R/R*) of all PCs in feedback, mismatch and playback phase, sorted by amplitude of mismatch response. None of the PCs are classified as nPE neurons (indicated by gray shading to the right). **c** Same as in (b) after plasticity. Somatic excitation and inhibition are balanced. PC population rate remains at baseline. All PCs classified as nPE neurons (also indicated by black shading to the right).

**Figure S4.**
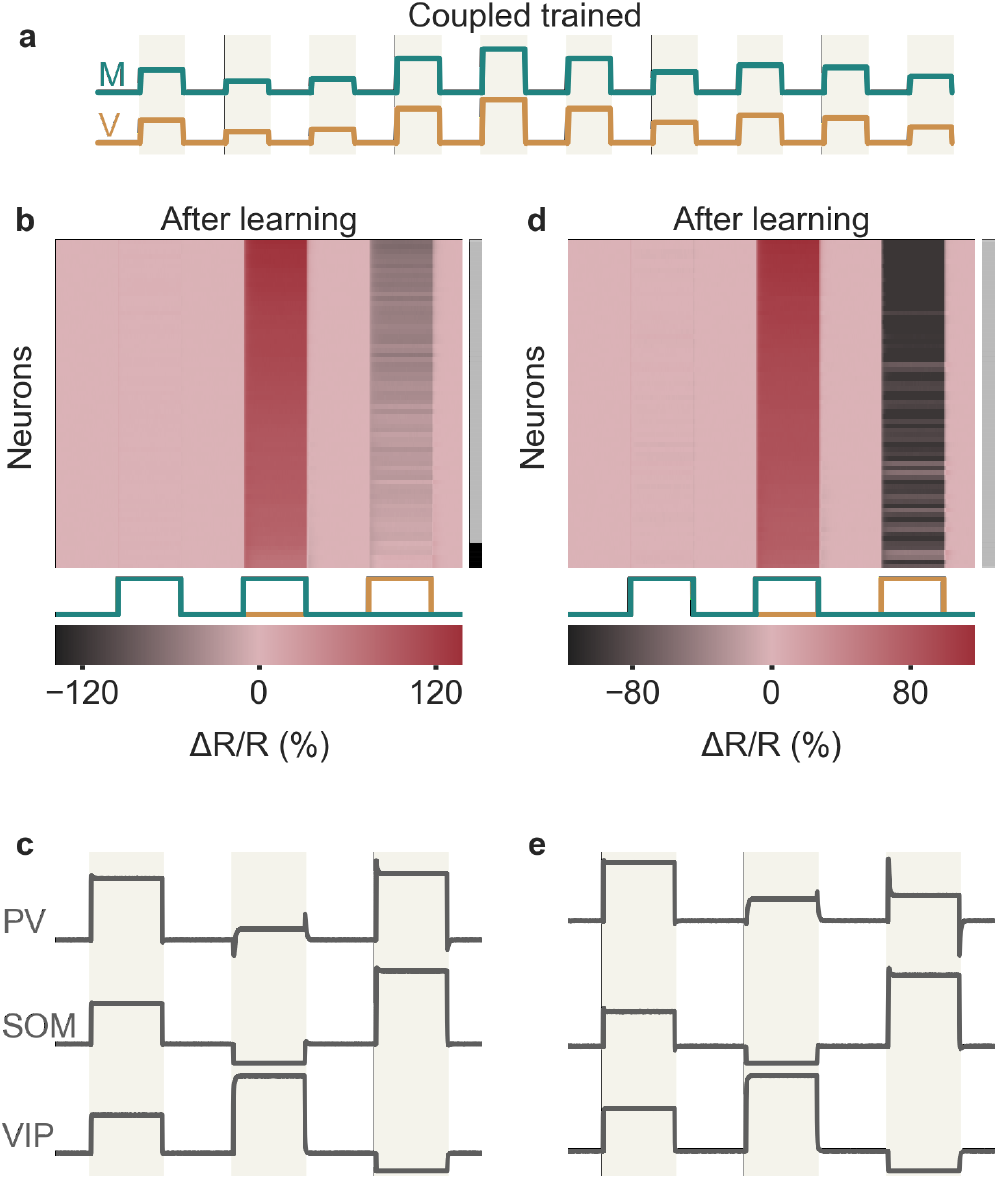
Coupled-trained networks can produce nPE neurons that decrease their activity in playback phase. **(a)** During plasticity, the network is exposed to a sequence of feedback phases only, representing perfectly coupled sensorimotor experience. Network model shown in Fig. 1. Connections from VIP to PV neurons are non-plastic. **(b-c)** Model in which an excess of dendritic inhibition does not affect the soma of PCs. Connection strength from VIP to PV neurons fixed to −0.3. **(b)** Response (Δ*R/R*) of all PCs in feedback, mismatch and playback phase, sorted by amplitude of mismatch response. All PCs increase their activity during mismatch phase but decrease their firing rate during playback phase. The decrease of PC activity during playback is a result of an excess of somatic inhibition mediated by PV neurons. **(c)** Population responses of PV, SOM and VIP neurons in all phases. Responses normalized between −1 and 1 such that baseline is zero. (d-e) Model in which an excess of dendritic inhibition is forwarded to the soma of PCs. Connections from VIP to PV neurons fixed at a value that ensures a balance of somatic excitation and somatic inhibition (see Eq. 34 in Methods). **(d)** Same as in (b). The decrease of PC activity during playback is a result of an excess of dendritic inhibition mediated by SOM neurons. **(e)** Same as in (c). PV neurons are less active during the playback phase than during the feedback phase.

**Figure S5.**
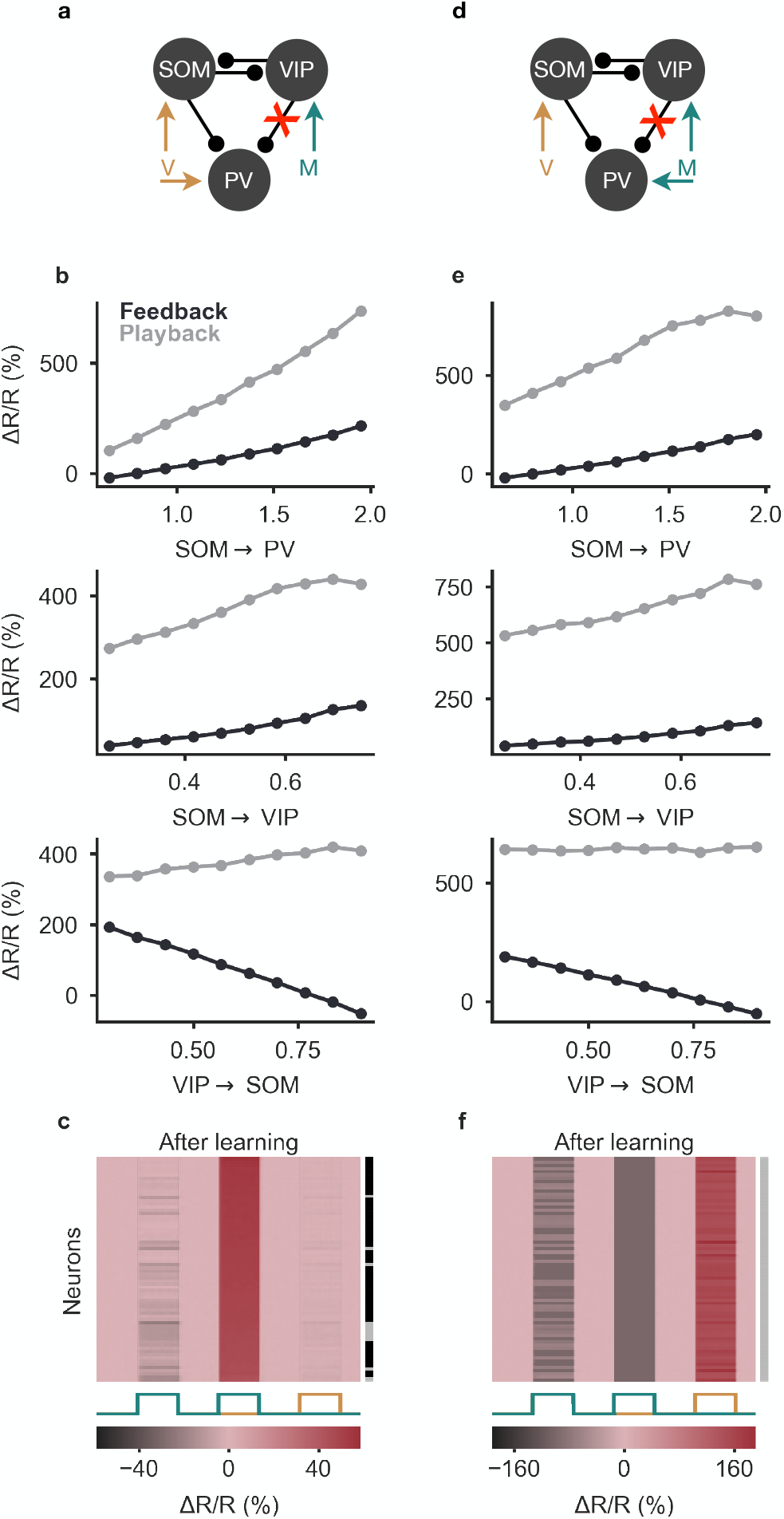
VIP→PV synapses are not required for the formation of nPE neurons. **(a)** Network model as in Fig. 1 but without VIP→PV synapses. PV neurons receive visual input. **(b)** Population response (Δ*R/R*) of PCs in feedback (dark gray) and playback phase (light gray) for varying SOM→PV (top), SOM→VIP (middle) and VIP→SOM (bottom) connections. For all values tested, firing rate during feedback and playback deviates from baseline. **(c)** Response (Δ*R/R*) of all PCs in feedback, mismatch and playback phase, sorted by amplitude of mismatch response. Most PCs change their firing rate only mildly in feedback and/or playback phase. As indicated by the gray/black shading to the right, many of the PCs are classified as nPE neurons. **(d)** Same as in (a) but PV neurons receive motor predictions. **(e)** Same as in (b) but PV neurons receive motor predictions. **(f)** Same as in (c) but PV neurons receive motor predictions. All PCs change their firing rate in response to all stimulation patterns. None of the PCs are classified as nPE neurons (indicated by gray shading to the right).

**Figure S6.**
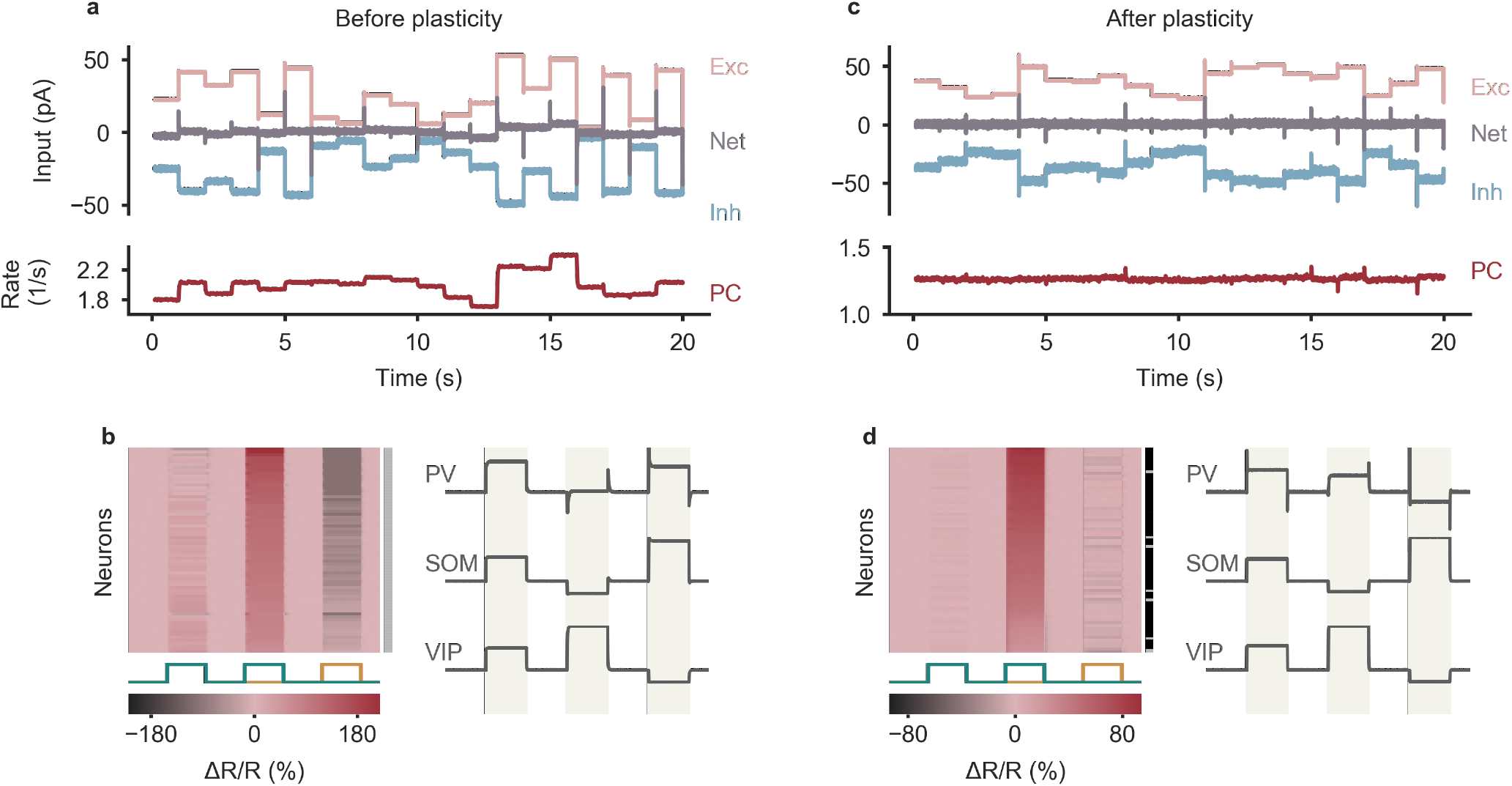
Balancing excitation, somatic and dendritic inhibition gives rise to nPE neurons in a model in which an excess of dendritic inhibition is forwarded to the soma. Network model, its inputs and the training set are shown in Fig. 1. Model setup modified to enable the presence of nPE neurons while abiding to Dale’s law: PCs receive 0.5 x visual input. External excitatory input onto the dendrites is set such that it balances inhibition mediated by SOM neurons in the baseline phase. Additional non-linearity for synapses from SOM neurons onto the apical dendrites of PCs: Δ*w*_DS_ ∝ *σ*(*A*_D_) · *A*_D_ · *r*_S_, where *A*_D_ denotes the total dendritic activity and *σ* is a sigmoid function. **(a)** Before plasticity, somatic excitation (light red) and inhibition (light blue) in PCs are not balanced. Excitatory and inhibitory currents are shifted by ± 20 pA for visualization. The varying net excitatory current (gray) causes the PC population rate to deviate from baseline. **(b)** Left: Response (Δ*R/R*) of all PCs in feedback, mismatch and playback phase, sorted by amplitude of mismatch response. All PCs change their firing rate in response to all stimulation patterns. None of the PCs are classified as nPE neurons (indicated by gray shading to the right). Right: Population responses of PV, SOM and VIP neurons in all phases. Responses are normalized between −1 and 1 such that baseline is zero. **(c)** Same as in (a) after plasticity. Somatic excitation and inhibition are balanced. PC population rate remains at baseline. **(d)** Same as in (b) after plasticity. Almost all PCs classified as nPE neurons (indicated by black/gray shading to the right). PV neurons are less active during the playback phase than during the feedback phase.

